# The concordance of length- and sequence-based STRs used in forensic markers with guidance for practice

**DOI:** 10.1101/2023.03.02.530748

**Authors:** Tikumphorn Sathirapatya, Wikanda Worrapitirungsi, Poonyapat Sukawutthiya, Hasnee Noh, Rachatipan Pitiwararom, Kornkiat Vongpaisarnsin

## Abstract

Next-generation sequencing (NGS) technology has shed light on every aspect of genetic discoveries, including forensic genetics. The Miseq® FGx Forensic Genetic System (Verogen) is one of the pioneering forensic NGS that provided a complete system from library preparation to data analysis. The system has been validated by several studies and led to a more practical aspect. Short tandem repeat (STR) is a well-established marker that was designed specifically for human individualization. Since NGS provides different data from fragment analysis, a new STR nomenclature is established to make NGS backward compatible with the previous data. In this study, Thai population were used to evaluate the Miseq® FGx Forensic genetic system (Verogen) in practical aspect, including concordance study and forensic population parameters. In summary, we purposed a practical guideline for sequence-based STRs.

## Introduction

It is undeniable that next-generation sequencing (NGS) technology will dominate in forensic DNA analysis. The Miseq® FGx Forensic Genetics System (Verogen) is one of the most well-known NGS systems designed specifically for forensic DNA analysis. The system provided a forensic test kit comprising various DNA markers that could increase the power of population statistics and the ability to genetic inference. In addition, this would have potential for forensic scenarios in a highly degraded situation or a low amount of DNA template that could be analyzed in a single reaction. For the Miseq® FGx system, the primary data was automatically analyzed by the built-in software of the sequencing system. Secondary analysis with managing the genetic data was performed with universal analysis software (UAS) (Verogen), which was designed exclusively for the Forenseq® Signature prep kit (Verogen). The software provides an easy-to-use interface and generates FASTQ files, performs alignment, and reports forensically relevant variants from NGS data. At present, this system has been thoroughly studied including the performance evaluations (1, 2), validation (3) and population studies (4–8). In 2016, the International Society of Forensic Genetics (ISFG) established guidelines for Short Tandem Repeat (STR) nomenclature (9) and guidelines for reporting population data (10) to facilitate the use of NGS in the forensic community. Unfortunately, some markers reported as reverse strand in UAS required additional modification to comply with ISFG guideline (7). However, both open source and commercial software were available as alternative analysis tools. Each software was based on a different way to extract the STR sequence from raw files and provided distinct advantages (11). For example, STRinNGS v2.0 was a free alternative software operated on Docker (12). This software provided a genotype report following ISFG guidelines and included flanking region sequences in the report. These flanking region sequences could assist in data interpretation, especially when a mutation or variation on the primer binding site was encountered (4).

It was a great challenge to transition to NGS technology without throwing away the previous technology. The novel STR sequence-based NGS should mutually benefit from a length-based STR (mostly from capillary electrophoresis), which was the largest database collected in the past. Both data should evaluate their compatibility and analyze the vagueness. In this study, Thai population were used to evaluate the concordance between CE length-based, UAS, and STRinNGS v2.0 for autosomal STR and Y-STR data generated from the Forenseq® Signature prep kit (Verogen). Furthermore, concordance genotypes were evaluated in forensic population statistics. Finally, the guidance was recommended for forensic practice.

## Materials and Methods

### Samples, Library Preparation, and Sequencing

Three hundred and ninety Thai individuals were included in this study. Samples were available in blood-impregnated FTA card, and all had length-based aSTR profiles from the Investigator IDplex kit (QIAGEN). A 1.2 mm. diameter FTA card was amplified with a DPMB primer mix of Forenseq® Signature prep kit (Verogen). Library preparation and sequencing were carried out according to the manufacturer’s instruction in 13 sequencing runs on Miseq FGx ^TM^ (Verogen). The quality metrics of each run (Cluster density, % cluster passing filter, % phasing, and % prephasing) and the total read count of each sample were included for further analysis (Fig 5.). This study was approved for ethical consideration by the Institutional Review Board of the Faculty of Medicine, Chulalongkorn University (IRB No. 104/64). The study was conducted retrospectively by using anonymous data.

### Data Analysis

In this study, only genotypes aSTR and Y-STR were used in data analysis. Initially, the sequencing data were analyzed by UAS version 1.3 (Verogen) with the default settings. The sample detail report.xlsx files were retrieved for aSTR and Y-STR genotypes. The sample inclusion criteria for performance and population genetic analysis are unrelated Thai individuals with a complete profile of aSTRs and YSTRs in the UAS report. All duplicate profiles were excluded (Fig 1).

Then, the FASTQ file of read 1 was reanalyzed using STRinNGS v2.0 (12) for allele calling of the aSTR and Y-STR markers. The configuration file was obtained from the supplement material of (12) which the allele nomenclature and designation were consistent with ISFG’s guideline (9).

**Fig 1.**
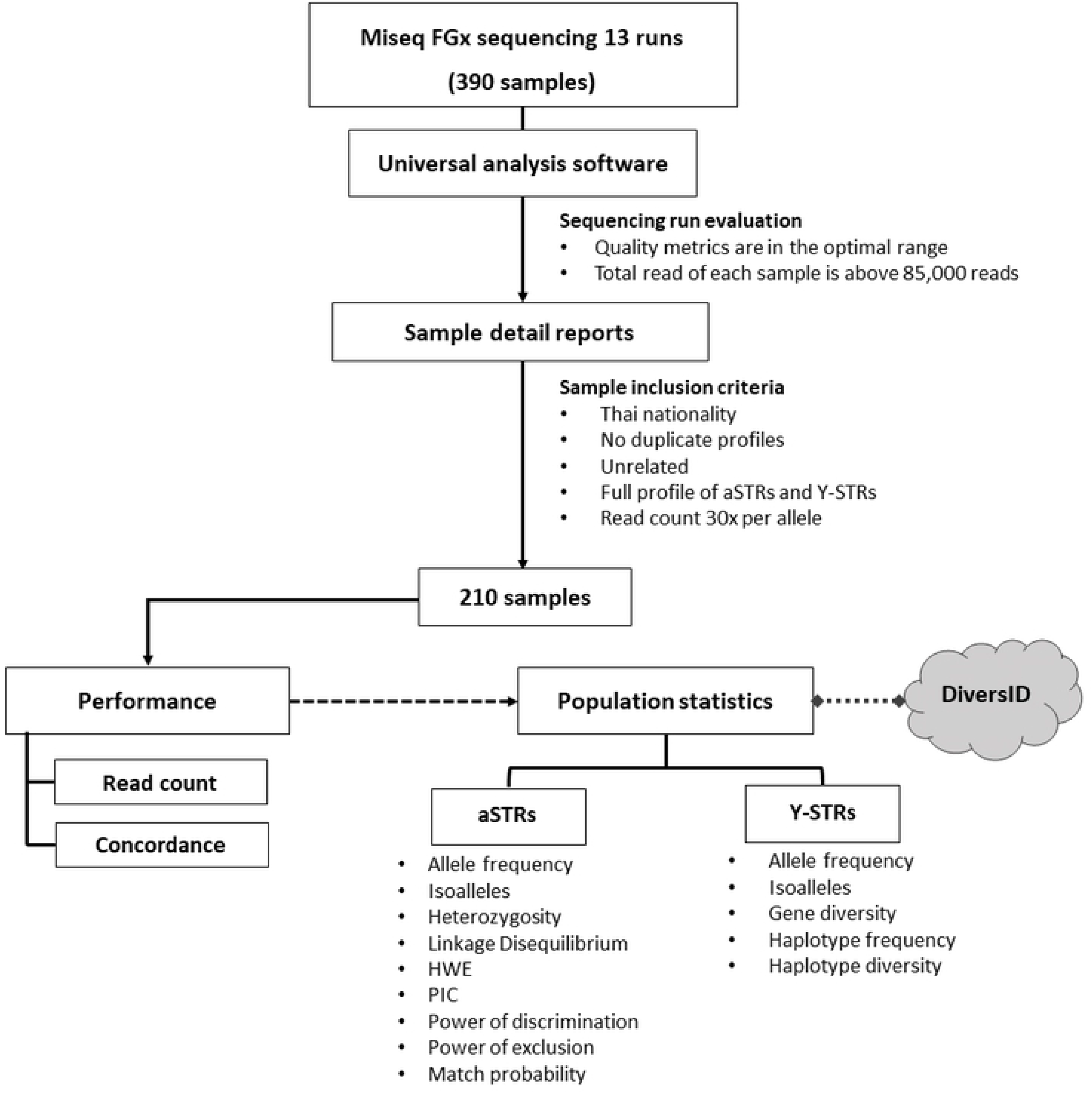
Workflow of the samples included for performance and population genetic analysis.

### Performance test

The performance or quality of the data was evaluated in two ways: the read count of alleles from the STRinNGS report and the concordance study between different analysis methods. Fifteen overlapping aSTR loci were compared for genotype concordance from three different analyzes: CE length-based, UAS, and STRinNGS. For the other 12 aSTRs and 24 Y-STRs, the concordance study relied on sequence-based genotypes between UAS and STRinNGS. The result was considered concordant if the reported genotype was identical in all analyses. If one of the reported genotypes differed from others, the disagreed genotype was investigated further and subdivided into discordant due to variation or allele designation, allele dropout, low read count sign, and stutter interference as a cause of disagreement. If the disagreed genotype was still inconclusive, the samples could be reamplified and reanalyzed by either Powerplex® Fusion 6C system (Promega) or Verifiler^TM^ amplification kit (Applied Biosystem) for aSTR and Yfiler™ plus PCR amplification kit (Applied Biosystem) for Y-STR. Then, the genotypes that accounted for allele dropout and stutter interference were manually edited, except discordant due to variation or allele designation was included without editing.

### Population statistics

Isoalleles are alleles with identical lengths in the base pair but contain different sequences including flanking region sequence in this study. An isoallele with the highest frequency is determined by its original allele number, and the others are assigned to an index number (S2.1, S2.3 and S2.4). Sequence-based data refer to sequencing data including isoalleles and length-based data indicated NGS data without isoalleles.

For aSTRs, the parameters of the forensic population that cover the number of alleles, allele frequency, expected heterozygosity (Hexp), the observed heterozygosity (Hobs), polymorphic information content (PIC), the power of discrimination (PD), power of exclusion (PE), the probability of match (PM), typical paternity index (TPI) and the Hardy-Weinberg equilibrium (HWE) were calculated by STRAF (30) (Fig 1). Furthermore, the combined PM and PE values were compared between the sequence-based data in this study and the DiversID data (https://diversid.info/). These data (N = 244) from DiversID were chosen to represent the length-based aSTR data since it was built from the UAS sample detail report and our included samples were a subset of this data set. The linkage disequilibrium (LD) was estimated with 10,000 permutations by Arlequin version 3.5.2.2 (15). For Y-STRs, allele frequency and gene diversity (GD) were calculated for each marker using GenAlex (31) and the GD formula from (32), respectively. The number of haplotypes and haplotype diversity were calculated by direct counting.

## Results

### Sequencing Run Evaluation

Thirteen sequencing runs demonstrated an average cluster density of 1,220.38 K/mm^2^ with an average cluster passing filter of 89.63%. The average phasing was 0.176% and the average prephasing was 0.282%. All average sequencing quality parameters were in the recommended range except the percentage of prephasing in all runs. This metric was constantly high (S1.1) in all sequencing runs. The percentage of prephasing is defined by unsynchronized strands that extended ahead of the current cycle in the cluster and resulted in less useable data output. The high percentage of prephasing can occur by the insufficient flushing in the flow cell or the incorporation of an ineffective 3’ terminator (13). Run no. 1 was the first sequencing run of this Miseq FGx^TM^ (Verogen) and showed both prephasing and phasing higher than optimal range (S1.1). Run no. 11 showed a cluster density of 1,797 K/mm^2^ and the cluster passing filter slightly fell from 80% (S1.1). This accounted for the overclustering.

### Performance

#### Read count

All samples from 13 sequencing runs were filtered according to the criteria in Fig 1. and resulted in 210 samples (129 males and 81 females) for concordance study and population genetic analysis. According to the STRinNGS genotyping, the highest average read count of aSTR was TH01 (7,827.90) and the lowest was D5S818 with an average read count of 245.19. For Y-STRs, the highest average read count was DYS438 (7,906.61) and the lowest was Y-GATA-4 (235.74) (S1.2). All included alleles showed a read count greater than 30x, except these eight alleles (S1.3), which were included with a read count in the range of 22 to 29 reads and the genotypes had been confirmed by length-based or UAS sequence-based genotypes.

#### Length- and sequence-based concordance

Fifteen overlapping aSTRs were compared between the Investigator IDplex kit (QIAGEN) and the Forenseq® Signature prep kit (Verogen) for length-based and sequence-based concordance. Genotype comparison was carried out between 3 different analyzes: length-based, sequence-based UAS, and sequence-based STRinNGS. Ten out of 15 aSTRs displayed complete concordant genotype (Fig 2) with the 99.56% average concordance. Most of the different genotypes were from STRinNGS and few genotypes were from UAS. D19S433 showed the least concordant locus that contained 96.2% length-based concordant genotype, 0.48% discordant, 2.86% dropout, and 0.48% stutter interference.

Two STRinNGS genotypes showed discordance due to variation or allele designation. Both discordant markers were not parsed in STRinNGS and showed ‘FALSE’ in the ExactParse column with two mismatch positions in the flanking region (up_or down_fl_num_diffs column) in the STRinNGS report (S1.4 and S1.5). The first discordant was found in CFS1PO where STRinNGS showed the 10,10.3 genotype, but length-based and UAS showed 10,11 genotype. The reported sequence of allele 10.3 with ATCT[10]atc showed two mismatch positions of the flanking region in STRinNGS and therefore disagreed with allele 11 in UAS and length-based (S1.4). The second was observed in the D19S433, where the STRinNGS genotype was 13,13.1, but the length-based and UAS genotypes were 13,13.2. Allele 13.1 with the repeat motif, reported sequence showed two mismatch positions in the 5’ flanking region in STRinNGS and thus disagreed with in UAS and length-based (1.5). Then, these two discordant results were reanalyzed using STRait Razor Online (SRO) (14). The SRO genotypes were conformed to UAS and length-based. However, repeats in bracket format were desolated and showed only allele number and sequence string (S1.4 and S1.5). The dropout occurred mainly in larger size alleles in most markers, which was found in D16S539, D5S818, and D19S433. STRinNGS reported the false homozygous genotype or complete dropout. For D16S539, UAS and STRinNGS showed heterozygous genotypes, while length-based genotypes showed severely heterozygous imbalanced genotypes. The heterozygous genotypes were confirmed by re-amplifying with the Verifiler ^TM^ amplification kit (Applied Biosystem). Stutter interference was found in D18S51 and D19S433 (Fig 2), where the read count of the stutter reached the interpretation threshold and was called by the STRinNGS.

**Fig 2.**
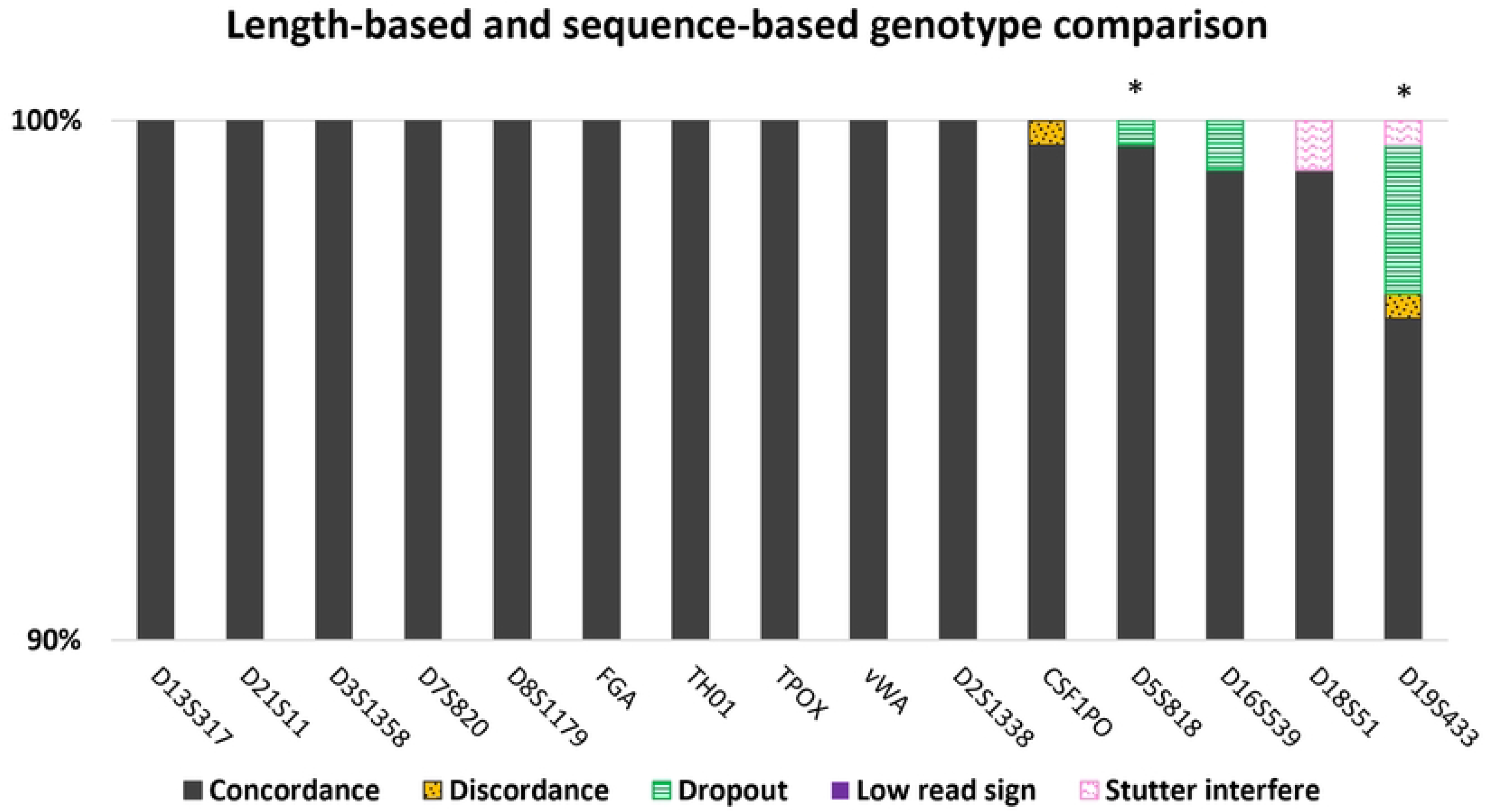
Comparison of length-based and sequence-based genotypes in 15 aSTRs. The gray bar represents the percentage of concordant. Color bars with pattern represent discordant result caused by discordant due to variations (yellow), allele dropout (green), low read sign (purple), and stutter interfere (pink) in STRinNGS. Asterisk (*) indicated that some of the UAS showed differences in genotype compared to STRinNGS and length-based.

#### Bioinformatic concordances

The comparison was divided into five categories (Fig 3). The result was counted as concordant if the reported genotype was identical in both analyses. If one of the reported genotypes differed from others, the disagreed genotype was further investigated and subdivided into discordance due to variation or allele designation, allele dropout, low read count sign, and stutter interference. Sixteen markers (eight aSTRs and 8 Y-STRs) of 35 markers were concordance (Fig 3) and the average concordant was 94.91%. DYS612 was the only Y-STR marker excluded from the concordance study due to the different allelic designation between length-based and the sequence based (4, 9). Discordant due to variation or allele designation category was not found among these 35 markers, and allele dropout was the primary cause that made the genotype different in STRinNGS results (Fig 3). The allele dropout category was found most in D22S1045, Penta E, DYS19, DYS392, DYS635, DYF387S1, DYS389II, Y-GATA-H4, DYS385, Penta D, DYS438, DYS448, and D10S1248, respectively. The low read sign was classified when the read count of the reported allele was low or showed ‘‘more than three alleles” in the PredAllele column in STRinNGS. This observation was found most in DYS19, DYS635, D22S1045, and DYS392, respectively. Lastly, stutter interference was scattered in most markers (Fig 2.) and encountered in DYS385, DYS635, DYS392, DYS389II, DYS437, DYS448, DYS481, DYS570, D22S1045, DYS390, DYS533, and DYS576. Since there was no CE length-based genotype available for these markers, some of these genotypes were reanalyzed by either the Powerplex® Fusion 6C system (Promega) or the Verifiler^TM^ amplification kit (Applied Biosystem) for aSTR and Yfiler^TM^ plus PCR amplification kit (Applied Biosystem) for Y-STR. The results that accounted for allele dropout and stutter interference were manually edited. Furthermore, the bi-allele at DYS533 and the tri and tetra-allele at DYS387S1 were excluded from population genetic analysis and were recorded separately in S1.6.

**Fig 3.**
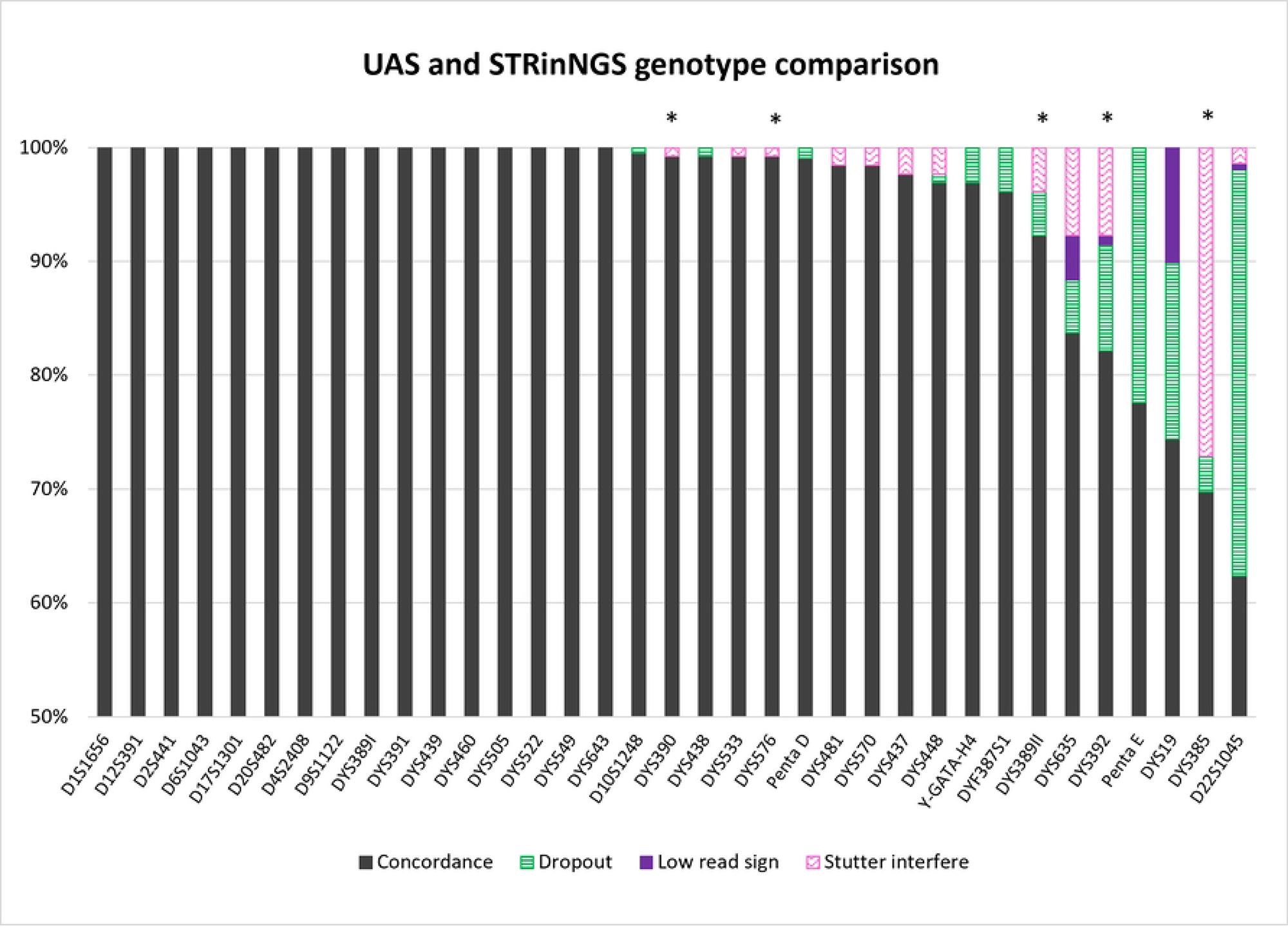
Comparison of the UAS and STRinNGS genotypes in 12 aSTRs and 23 Y-STRs. The gray bar represents the percentage of concordant. Color bars with pattern represent discordant result caused by allele dropout (green), low read sign (purple), and stutter interfere (pink) in STRinNGS. The asterisk (*) indicated that some of the UAS showed differences in genotype compared to STRinNGS or length-based.

#### Isoalleles and flanking region variants

After samples were filtered and examined for concordance, the population genetic analysis relied on data from the STRinNGS report that included allele number, repeat sequence, and flanking region sequence. Isoalleles and microvariants were assigned a new allele number that referred to their original allele number (S2.1, S2.3 and S2.4). In Fig 4, the observed alleles of each marker were subdivided into the length-based allele (LB), the number of alleles gained by variation in the repeat region (RR) and the number of alleles gained by variation in the flanking region (FR) for aSTR (Fig 4a) and for Y-STRs (Fig 4b). The heterozygosity and gene diversity values were plotted against the number of alleles of aSTR and Y-STR, respectively. More than half of the STRs (62.7%) gained more alleles from sequence-based data. These isoalleles were observed in the compound or complex repeat STR group more than in the simple repeat STR group.

**Fig 4.**
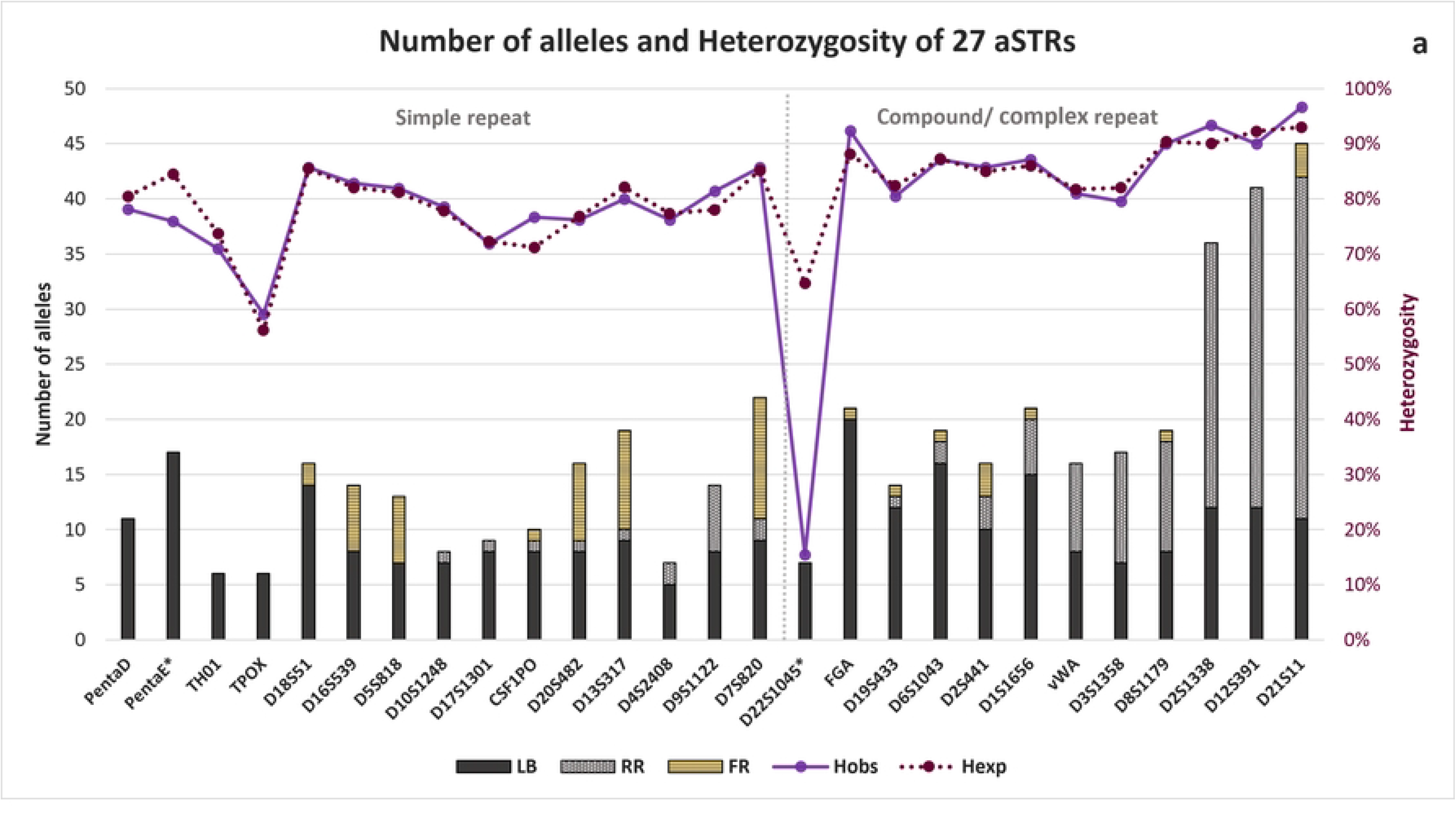
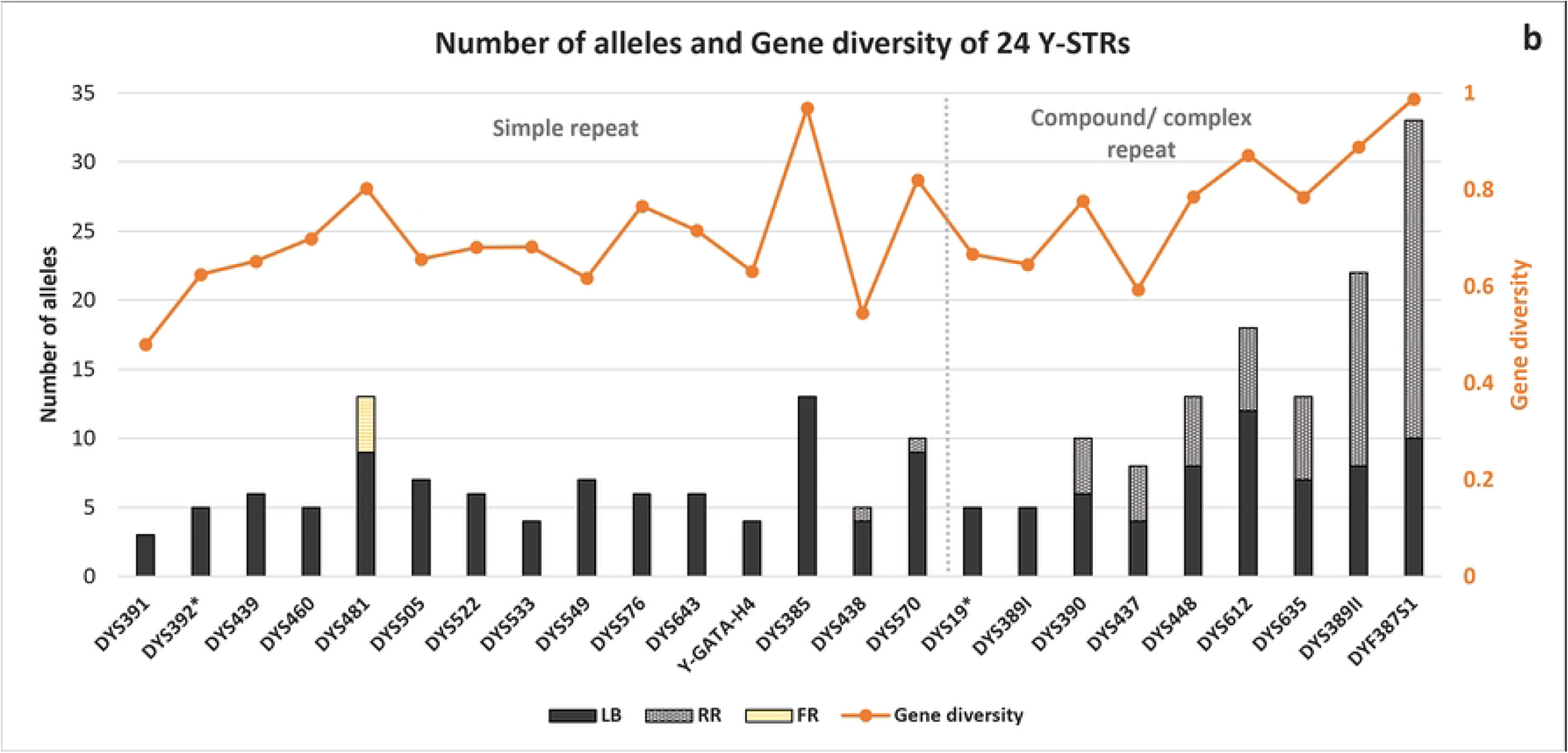
Number of alleles, heterozygosity, and gene diversity in aSTR (a) and Y-STR (b), respectively. LB stands for the length-based allele. RR indicated an allele gained by variation in repeat region and FR was an allele gained by variation in flanking region. The grey dashed line divides the simple repeat STRs from compound or complex repeat STRs. The asterisk (*) indicates a poor performance marker.

All rare variants (frequency less than 0.01) were recorded separately in S2.9. These rare variants were observed by the number of mismatched nucleotides in the flanking region (up_ and down_fl_num_diffs columns) and did not be determined in the rs_info of STRinNGS’s configuration file. The mismatched nucleotide was then identified and compared with the dbSNP database according to its nucleotide position.

#### Autosomal STRs

As compound and complex repeat aSTRs were made up of more than one repeat motif, they tended to share more isoalleles compared to simple repeat aSTRs (Fig 4a). The highest observed number of alleles was D21S11 (45 alleles), composed of the allele gained by RR (68.8%), the allele gained by FR (6.6%) and the length-based allele (24.4%). The simple repeat STR group gained alleles by the flanking region variation except in D9S1122, D4S2408, D17S1301 and D10S1248; the number of alleles increased only by RR. Five aSTRs (PentaD, PentaE, TH01, TPOX, and D22S1045) did not show additional alleles compared to length-based alleles. D7S820 was the highest number of alleles obtained by variation in the flanking region (11 alleles). These came from the combination of two SNPs located on each side of the repeat region of D7S820. These were rs7789995 (A>T) on the 5’ flanking region and rs16887642 (G>A) on the 3’ flanking region (S2.1). The expected and observed heterozygosity showed similar values among 27 aSTRs, except for D22S1045, which was marked as a poor performance marker (Fig 4a).

#### Y-chromosomal STRs

DYS387S1 showed the highest number of alleles (33 alleles) (Fig 4b). Nine out of 24 Y-STRs gained more alleles by only RR that were DYS387S1, DYS389II, DYS635, DYS612, DYS448, DYS437, DYS390, DYS570, and DYS438. A simple repeat Y-STR, DYS481, was the only marker that gained more alleles by the combination of two SNPs in the flanking regions. There are rs368663163 (G>A) and rs370750300 (G>T) located in the 5’ flanking region (S2.3). The gene diversity of 24 Y-STRs was in the range of 0.4797 - 0.9875. The highest value of gene diversity was DYS387S1 and the lowest was DYS391.

### Population statistics

#### Autosomal STRs

Allele frequencies, allelic range, and STR repeat region sequence of each aSTR were displayed in S2.1. Population statistical parameters were evaluated for 27 sequence-based aSTRs (S2.2.1). All loci were analyzed based on the total number of 420 alleles except D22S1045 (N = 336) and PentaE (N = 366). These two markers suffered from dropout of larger alleles and, therefore, were excluded from population statistics analysis. The expected heterozygosity (Hexp) was greater than 0.7000 (0.7120 - 0.9300), except TPOX (0.5615). The highest heterozygosity was D21S11. The polymorphic informative content (PIC) was in the range of 0.5080 - 0.9235. The highest PIC was D21S11 and the lowest was TPOX. For the typical paternity index (TPI) value, 25 STRs were found in the range of 1.2209 - 15.0000 from TPOX and D21S11, respectively. The more observed allele resulted in an increase in PIC and TPI values, such as D21S11, which showed the highest number of observed alleles, and showed the highest PIC and TPI values.

When comparing the combined value of PM and PE between sequence-based data and length-based data, the combined PM was decreased approximately 3,000 times when considering sequence-based data. Furthermore, the combined PE of the sequence-based data was higher almost 18 times compared to combined PE from DiversID (S2.2.2). The 25 markers were in Hardy-Weinberg equilibrium (HWE) after Bonferroni correction (p-value > 0.001). After testing for linkage disequilibrium using Arlequin (15), a pair of D7S820 and D3S1358 was significant (p-value > 0.001) regardless of Bonferroni correction (S2.5).

#### Y-chromosomal STRs

The allele frequency of each marker was in S2.3 and S2.4 for biallele Y-STRs. All loci were analyzed based on the total number of at least 120 alleles, except DYS19 (N = 97) and DYS392 (N = 52). These two markers showed a low read count in most of the sample and resulted in a few numbers of analyzable alleles. Therefore, DYS19 and DYS392 were excluded from further analysis. The gene diversity (GD) was in the range of 0.4796 - 0.9875. The highest GD was shown in DYS387S1 (0.9875) and the rest of the Y-STR showed GD higher than 0.5000 except DYS391 (Fig. 4b and S2.6). From a total of 20 Y-STRs, 117 unique haplotypes were observed from 119 samples with one common haplotype (S2.7).

## Discussion

The road to implementing NGS technology is near and shows a promising future for forensic DNA analysis. An important milestone is establishing population data for STR markers (16). These population data were essentially built from high-quality and reliable sequencing data since these data will be applied for statistical probability or to calculate the strength of DNA evidence.

Sequencing quality metrics for the illumina platform included cluster density, percentage of cluster passing filter, phasing, and pre-phasing. Overclustering was one of the common challenges when the user validated or altered the library preparation protocol. The impact of overclustering can be various. If the overclustering is over-crowded, this may result in complete run failure. On the other hand, if the cluster density was slightly above the maximum, the % cluster passing filter decreased and consequently there were fewer sequencing data. According to Forenseq’s manufacture instruction, bead-based normalization was already included, and 7-μl of pooled library was specified for loading. Library quantification was not needed. Our wide range of cluster density (899-1,797 K/mm^2^) was similar to (17). Many factors can influence the cluster density including number of samples per run, library normalization, library quantification, and pooled library concentration (18). We recommended conducting the quantification of the qPCR library and adjusted the amount of pooled library prior to sequencing. Moreover, problems with phasing and prephasing can be encountered. These quality metrics are related to the fluidic system and the effectiveness of a reversible terminator. As phasing and prephasing created noise in the overall signal of cluster, these metrics were assumed to increase sequencing error rates (13). We found a constant high percentage prephasing even though post-run wash and maintenance wash were carried out according to the Miseq user guide. Therefore, to our knowledge, maintenance of the fluidic system was crucial and can improve the quality of sequencing data. Samples with a total read count of less than 85,000 reads should be processed with caution. Both sequencing quality metrics and total read count were effective parameters to evaluate the overall quality of sequencing data. Then, a sample detail report of each sample was used to screen the STR profile with criteria as shown in Fig 1 and resulted in 210 samples. In addition, we experienced a consistent low total read count of the 2800M positive control. This complied with the consistent incomplete profile of 2800M in (19) and therefore we recommended increasing the DNA input from 1 ng to 5 ng (data not shown).

Two different analysis tools were used for genotyping STR of sequencing data. The first one is UAS, a default analysis tool that automatically analyzes the sequencing data from the Miseq FGx machine. UAS provided a user-friendly interface with a default analysis setting that was suitable for both amateur and professional users (3). The alternative analysis tool was STRinNGS v.2 (12). This open-access analysis tool required command-line skills. The advantage of STRinNGS was the configuration file (9) that has been validated for Forenseq’s library and designed to interpret data according to ISFG’s recommendation (9). In version 2, the STRinNGS report concentrated on the similarity of the flanking region between the sample and the human reference genome. According to the read count (S1.2), the broad range of read count in STRs was expected as the Forenseq® Signature prep kit (Verogen) relied on large multiplex PCR. The efficiency of each marker in multiplex PCR was different and affected the amount of read count (2). UAS demonstrated that the read count of each marker differed from STRinNGS (data not shown). However, both tools provided most of the allele with a read count higher than 30x. The different read count was assumed to be due to the different lengths of the analyzing sequence string. These have been mentioned in (5) where the read count was compared between STRait Razor and UAS.

For the concordance study, allele calling of different analysis tools was compared to evaluate similarity and the backward compatibility to the conventional technique. Length-based and sequence-based concordance studies demonstrated high concordance (99.56%) in 15 aSTRs (Fig 2). Among the five categories used for classifying the comparison, allele dropout, low read count sign, and stutter interference were easily recognized and editable except the discordant due to variation or allele designation. This category was observed in two genotypes from two different markers (S1.4 and S1.5). Both genotypes showed mismatch points in the flanking regions. A discordant allele was found in the STRinNGS report compared to length-based and sequence-based UAS results. The flanking region that was adjacent to the repeat region displayed identically to the repeat motif sequence. This appearance made the boundary of the repeat region ambiguous. The unknown insertion/deletion that potentially disrupted the boundary of the flanking region was suspected to be the reason for these discordant results. This category had potentially produced a conflict of interpretation between the length- and sequence-based data. Therefore, previous studies (4, 6) suggested including flanking region sequence along with the reported allele. As the STRinNGS algorism relied on an approximate match of the flanking region sequence, this analysis tool would be more susceptible to these variations. In this study, we included these two discordant alleles with their flanking region sequences in the population data analysis.

Furthermore, the bioinformatic concordance between UAS and STRinNGS in the other 35 STRs resulted in 94.91% average concordance. In Fig 2, dropout, stutter interference and low read count sign categories were the main cause for disagreed results between these two analysis tools. D22S1045 and DYS392 were well-known markers of poor performance in Forenseq® Signature prep kit (Verogen) (3, 7, 20, 21) and a high percentage of dropout was expected. Moreover, in this study a high percentage of dropouts was first reported in Penta E and DYS19. It should be noted that a high percentage of stutter interference in DYS385 comes from UAS results (data not shown). Stutters usually were allele positioned between small and large alleles of the heterozygous genotype regardless of their read count (data was not shown). These occurred only in an older version of UAS, as the updated version was not found. Overall, these disagreed results presumably originated from the different analysis settings between UAS and STRinNGS. The minimum read count for genotype call in STRinNGS was equal to or more than 100 (default setting) (12), making STRinNGS more stringent than UAS. However, the more stringent the analysis setting was, the more data was lost. Therefore, some markers may require additional optimization for the STRinNGS analysis setting. In this study, we focus on concordance and population data for forensic DNA analysis and leave this issue for our future study. The disagreed genotypes were reanalyzed to confirm the results using one of the CE commercial STR kits and manually edited prior to population data analysis. Therefore, CE length-based data was crucial to evaluate the concordance of bioinformatic analysis.

Unfortunately, we used retrospective CE length-based data, resulting in a limited number of comparable STRs. As commercial NGS kits contained more markers, none of the recent commercial CE kits can share 100% comparable markers. The Powerplex® Fusion 6C system (Promega) or Verifiler^TM^ amplification kit (Applied Biosystem) was the superior choice and provided 23 comparable aSTR. For Y- STRs, we recommended the Powerplex Y23 system (Promega) or the Yfiler^TM^ Plus amplification kit (Applied Biosystem) that contained 19 comparable Y-STRs. Furthermore, two or more different analysis tools were recommended for forensic sequencing data to minimize errors in data interpretation (11).

The STRinNGS configuration file from supplement materials (12) has already specified the most common variants in the flanking region that can be found in most of the population. Most STRs were increased in the number of alleles compared to the number of length-based data and were encountered in aSTRs more frequently than Y-STRs (Fig 4). The variation in the repeat region was the main cause of increasing alleles and was more frequent in the compound/ complex repeat STR group. These findings were consistent with other studies (4, 8, 22). Four markers (Penta D, Penta E, TH01, and TPOX) that did not receive additional information from the sequencing data in this study were shared among the other studies (6, 22, 23). Some markers such as D16S539 and D20S482 that gained isoalleles by variation in the flanking region can become no isoalleles when considered in the repeat region sequence alone. For UAS sample detail report, only repeat region sequence was shown, and thus variation in the flanking region became invisible. In contrast, D18S51 was one of aSTR that the UAS sample detail report displayed a repeat region with 10 bp of the 3’ flanking region. This 10 bp of the 3’ flanking region covered rs535823682 (A>G) and thus became easy to access for UAS’s user. Variations in both FR and RR of D22S1045 were reported by several studies (4, 24, 25) and were also excluded from population data due to its poor performance. These indicated that the poor performance of this marker may camouflage the discovery of variations in FR and RR. DYF387S1 was responsible for rapid mutating Y-STRs and gained the most advantage from the sequencing data. The complexity of its repeat motif tended to generate more isoalleles. This finding was in agreement with other studies (4, 23, 26, 27). Many Y-STR markers were reported with variation in FR (26) where in our study were not; DYF387S1, DYS389I, DYF389II, DYS385, DYS612, DYS390, DYS437, DYS438, DYS576, DYS522, DYS460 and Y-GATA-H4. This may be due to our small sample size (N = 129). The variations in DYS19 and DYS392 were influenced by its poor performance similar to D22S1045. It is interesting that DYS481 can be classified as compound/complex repeats Y-STRs group (26). This marker is composed of a simple repeat of CTT with two SNPs (rs368663163 and rs370750300) in the 5’ flanking region. rs370750300 located next to the beginning of the repeat region. When considered 3-bp including rs370750300 (CTG > CTT) as an uncounted repeat pattern variation, the repeat region pattern became [CTG]_0-2_ [CTT]_n_.

According to (16), ambiguous results can be introduced in many ways when comparing DNA sequences between data sets. Different genomic interval that included for the secondary analysis tool was one of the reasons, even though from the same library preparation kit. Analyzable regions were used for main forensic commercial kits, including the Forenseq® Signature prep kit (Verogen) to facilitate the implementation of NGS in forensic fields (16, 28). The STRinNGS v2.0 (12) was extended the analyzable regions and may require editing to fit the analyzable regions.

We did not report novel variants in repeat and flanking region sequences with a frequency of more than 1%. Lastly, all observed variants in the flanking region were compared with other populations (S2.8). Four SNPs from four different STRs in this study showed different MAF compared with others; rs73801920 in D5S818 (A: 0.1214), rs16887642 in D7S820 (A: 0.2452), rs7460515 in D2S441 (A: 0.081) and rs75219269 in vWA (G: 0.2595).

Some variations discovered in this study were noteworthy. D21S11, D12S391, and D2S1338 were alleles obtained that were consistent much more than other STRs regardless of the different secondary analysis tools used (4, 8, 22, 23, 25, 27). These aSTRs were compound/complex tetranucleotide repeats and gained the majority of isoalleles by their own variable in the number of repeat motif. Therefore, analysis tools based on different flanking region sequence boundaries would show insignificant different results compared to other studies. According to (29), common repeat motif of a D21S11 consisted of variable number of repeat blocks of TCTA and TCTG and surrounded by 43 bp of TCTA [3] TA [1] TCTA [3] TCA [1] TCTA [2] TCCATA [1] or 39 bp of TCTA[3] TA[1] <u>TCTA[2]</u> TCA[1] TCTA[2] TCCATA[1], respectively. The underline letter indicated the difference between these 43 bp sequence block in the repeat region. We observed 47 bp of TCTA [3] TA [1] TCTA [4] TCA [1] TCTA [2] TCCATA [1] within allele 32.2 with a frequency of 0.1119. This allele 32.2 was in accordance with (24) (GenBank: MH174777.1) with frequency of 0.0103 in the Asian population. Furthermore, rs75219269[G] was observed only in allele 14 of vWA with the sequence of TAGA [3] TGGA [1] TAGA [3] CAGA [4] TAGA [1] CAGA [1] TAGA [1] and the frequency was 0.2595 (S2.1). Compared to other studies, this allele 14 was found most in Asian population such as East Asian (0.2456) (4), Korean (0.1928) (23), Taiwanese (0.3151) (25), Tibetan (0.1916) (22) and Uyghur population (0.1269) (27).

For population statistics, we choose STRAF (30) for analysis. It was an open-access online tool that was easy to use and could provide all essential population statistics parameters, including the linkage disequilibrium. However, according to (30), the authors suggested performing LD with other software since STRAF could not provide a permutation test. Therefore, we selected Arlequin (15) for LD with 10,000 permutations. The 25 aSTRs were in Hardy-Weinberg equilibrium after Bonferroni correction (p-value > 0.001). A pair of D7S820 and D3S1358 was significant (p-value > 0.001) when LD was performed.

Finally, we created the study of the practical guideline for the concordance of sequence-based STR data (Fig 5). Secondary analysis with alternative software should be performed to confirm genotype. In case of discordance observed in any marker, length-based was recommended to determine the cause, which possible be from mutation or poor performance of the marker with sequencing-based study.

**Fig 5.**
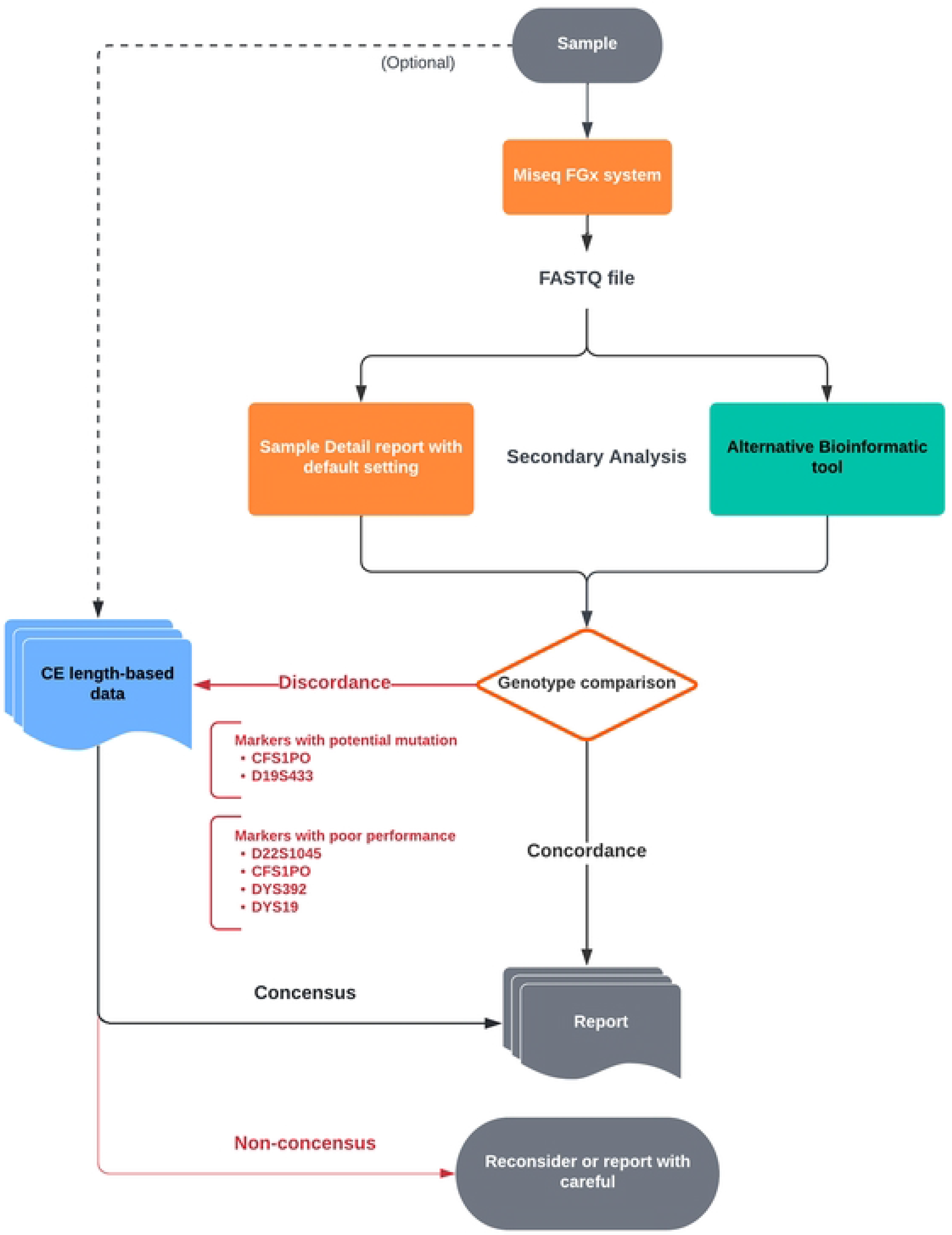
Practical guideline for sequence-based STRs.

## Acknowledgments

We would like to thank our colleagues from Forensic Genetics Research Unit, Department of Forensic Medicine, Faculty of Medicine, Chulalongkorn University, who provided insight and expertise that greatly assisted this research.

## Supporting information

**Supplement Table 1** composed of

**Supplement Table 1.1** Quality metrics of 13 pooled library sequencing runs.
**Supplement Table 1.2** Read count per locus of aSTRs and Y- STRs.
**Supplement Table 1.3** Eight alleles displayed less than 30 reads.
**Supplement Table 1.4** A discordant result of CFS1PO.
**Supplement Table 1.5** A discordant result of D19S433.
**Supplement Table 1.6** Observed bi-allele, tri-allele and tetra-allele in Y-STR genotype.

**Supplement Table 2** composed of

**Supplement Table 2.1** Allele frequency of 27 aSTRs.
**Supplement Table 2.2** composed of

**Supplement Table 2.2.1** Population statistics of 27 sequence-based autosomal STRs.
**Supplement Table 2.2.2** Comparison of combined PM and PE values between 25 sequence- based and length-based DiversID aSTR data.
**Supplement Table 2.3** Allele frequency of 22 Y-STRs.
**Supplement Table 2.4** Genotype frequency of DYF387S1 and DYS385.
**Supplement Table 2.5** Significant Linkage disequlibrium of 27 aSTRs using Arlequin.
**Supplement Table 2.6** Gene diversity (GD) of 24 Y-STRs.
**Supplement Table 2.7** Haplotype frequency and haplotype diversity of 20 Y- STRs.
**Supplement Table 2.8** Allelic frequency of flanking region variant in STRs.
**Supplement Table 2.9** Rare observed flanking region variants.

